# Enhanced-Sampling Molecular Dynamics Recovers Rare Functional RNA Conformations Across Diverse Structural Contexts

**DOI:** 10.64898/2026.08.22.746453

**Authors:** Deng Li, Megan Ken

## Abstract

Accurate determination of RNA conformational ensembles is essential for understanding RNA function and advancing RNA-targeted drug discovery, yet lowly-populated alternative states remain difficult to resolve with atomistic detail. A central constraint is that experimental refinement can only select conformations already present in the starting library, making library generation the limiting step. Using the HIV-1 trans-activation response element (TAR) as a model system, we benchmarked conventional MD (cMD) against the enhanced sampling methods Gaussian-accelerated MD (GaMD), replica-exchange Gaussian-accelerated MD (Rex-GaMD), replica-exchange with solute tempering (REST2), and temperature replica-exchange MD (T-REMD), as well as the structure-prediction based methods FARFAR2 and AlphaFold 3. Each library was refined against experimental residual dipolar couplings (RDC) and validated independently using ensemble-averaged QM/MM chemical shifts. We showed that T-REMD produced the most accurate ensemble by both measures, and its advantage tracked with broader, more continuous coverage of the interhelical conformational landscape. Broad temperature-range T-REMD also sampled conformations resembling excited state 1 (ES1) and the U23-A27-U38 base-triple, without requiring these states to be specified during library generation. More accurate ensembles further improved coverage of experimentally observed ligand-bound TAR conformations and enhanced ensemble-based virtual screening, linking structural accuracy to functional utility. The same workflow applied to the preQ_1_ class I riboswitch and the UUCG tetraloop improved agreement with experimental data in both cases. Together, these results establish replica-exchange enhanced sampling, particularly T-REMD, as an effective strategy for constructing experimentally validated RNA ensembles and accessing conformations corresponding to rare functional substates.

**Graphical Abstract:** 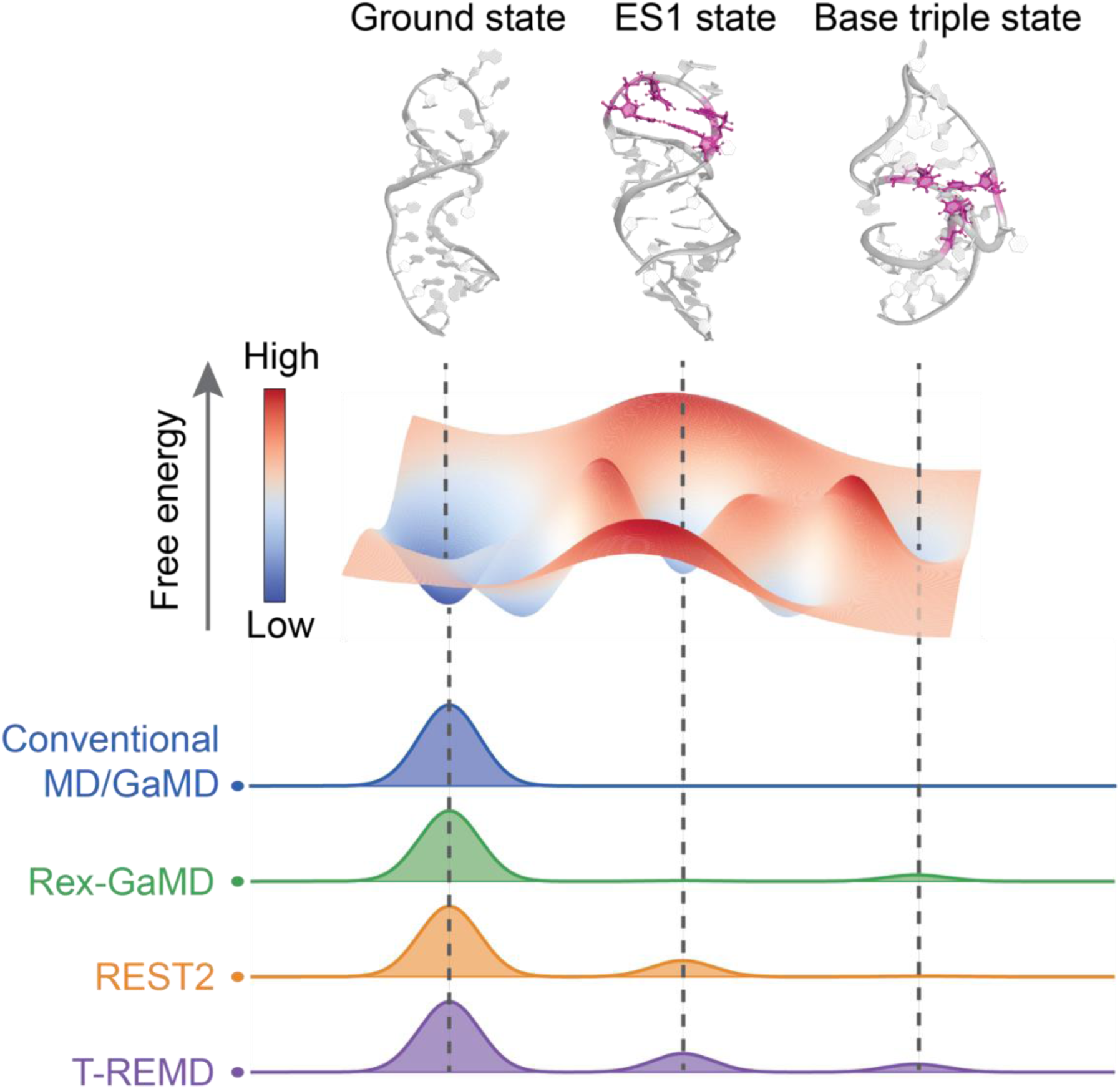

## Introduction

While the importance of RNAs was first understood as a genetic messenger relaying sequence information^1^, we now know that RNAs also carry out essential cellular functions by folding into 3D structures that can then form selective interfaces for proteins and small molecule recognition^2–4^. These recognition events play central roles in functions such as gene regulation^5,6^, transcriptional and translational control^5,7^, environment sensing^6^, and scaffolding ribonucleoprotein assemblies^2,3^. These functions are often not encoded by a single folded structure, but by the ability of an RNA to redistribute among an ensemble of alternative conformations in response to its molecular environment^8^. As such, RNA-mediated processes are governed by an underlying conformational landscape, where dominant states as well as low-populated states and transient structural rearrangements can determine recognition, regulation, and catalysis^2,9–14^. Because of this, the ability to accurately characterize these landscapes is valuable, allowing for prediction of cellular activity^15,16^, identification of drug targets^17,18^, and discovery of RNA-targeted therapeutics^19^. Experimentally defining these states at atomic resolution remains challenging. The most accurate way to measure conformational dynamics is using nuclear magnetic resonance (NMR), which is powerful because it is highly sensitive to minute dynamical changes on several timescales^20^. However, most NMR observables report on ensemble-averaged properties, whereas the number of structures and population weights required to describe an RNA free-energy landscape far exceeds the information content of the measurements that can typically be acquired. Thus, even when experimental data reveal conformational heterogeneity, converting these data into a physically realistic atomistic ensemble remains fundamentally underdetermined^2,10–13^.

Computational modeling provides the atomistic hypotheses needed to interpret ensemble-averaged measurements and connect RNA structure to function. Approaches fall broadly into physics-based methods, which use atomistic or coarse-grained force fields to explore conformational space^21,22^; knowledge-based methods such as Fragment Assembly of RNA with Full Atom Refinement (FARFAR), which assembles structures from fragments drawn from experimentally determined RNAs and refines them with an all-atom scoring function^23^; and machine-learning methods such as AlphaFold 3, which infer atomic structures from patterns learned from sequence and structural data^24^. Among physics-based approaches, molecular dynamics (MD) simulations have been the favored methodology for sampling energetic landscapes as they can link local rearrangements to larger-scale collective motions, generate candidate conformers for experimental refinement, and provide mechanistic models of conformational exchange^25,26^.

However, the quality of an MD-derived conformational library depends on both the nucleic acid force field and solvent model, as RNA conformational equilibria are highly sensitive to stacking, hydrogen bonding, ion interactions, and solvation^27–29^. Accurate RNA simulations therefore require a balanced force field and solvent model combination rather than an optimized force field alone^30^. Even with improved models such as the four-point OPC water model^31^, conventional MD (cMD) often incompletely samples low-population conformations that can strongly influence experimental observables and molecular recognition.

Much of the work in RNA ensemble determination has been possible because of extensive structural information that is available for a specific viral RNA, the HIV-1 trans-activating response element (TAR)^32–35^. TAR consists of a helix-junction-helix motif that undergoes conformational changes to recognize the viral protein Tat and initiate transcription of the viral genome^33,34,36^. Early studies of TAR combined cMD with NMR residual dipolar couplings (RDCs) and showed that this strategy can recover RNA dynamical ensembles and reveal how local bulge motions are coupled to global interhelical rearrangements involved in adaptive protein recognition^25,37^. These RDC-guided ensemble studies have also shown that broader conformational libraries produce more accurate experimentally refined ensembles, underscoring a simple but important constraint that refinement can only select states that are already present in the starting library^25,38^. Increasing the breadth and representativeness of the conformational library is therefore central to improving RNA ensemble determination.

Structure-prediction approaches such as FARFAR-NMR address sampling by generating RNA conformational libraries from predefined secondary structures, followed by RDC-guided selection and QM/MM chemical-shift validation^23,30^. Applied to HIV-1 TAR, FARFAR-NMR improved ground-state ensemble accuracy relative to an cMD-derived library, particularly in the flexible bulge^30^. However, TAR also transiently populates ES1 (∼13%), ES2 (∼0.4%), and a rare U23–A27–U38 base-triple conformation involved in Tat recognition^11,13,36^. An atomistic model of ES1 was subsequently constructed by using relaxation-dispersion-derived secondary-structure information to generate separate ground- and excited-state libraries, which were jointly optimized against RDCs and validated using chemical shifts and state populations^14^. Although these studies established a powerful route for modeling low-population RNA states, they leave open whether such functional conformations can be sampled directly from physically generated trajectories without specifying their secondary structures in advance^9,14,30,38^.

Here, we test whether enhanced-sampling MD can generate broader and more representative RNA conformational libraries than cMD, improving experimentally guided RNA ensemble construction by sampling alternative states without having to specify them *a priori* (Figure 1). In cMD, a single trajectory is propagated at a fixed temperature on the unmodified force-field potential, and transitions between metastable RNA conformations may remain infrequent when the states are separated by high free-energy barriers. Enhanced-sampling methods promote barrier crossing either by modifying the potential-energy surface or by coupling simulations performed under different thermodynamic or Hamiltonian conditions. Using HIV-1 TAR, we benchmark MD, Gaussian-accelerated MD (GaMD), replica-exchange Gaussian-accelerated MD (Rex-GaMD), REST2, and temperature replica-exchange MD (T-REMD) for conformational library generation, and further test three T-REMD temperature windows to directly evaluate how sampling breadth affects ensemble quality.

**Figure 1.**
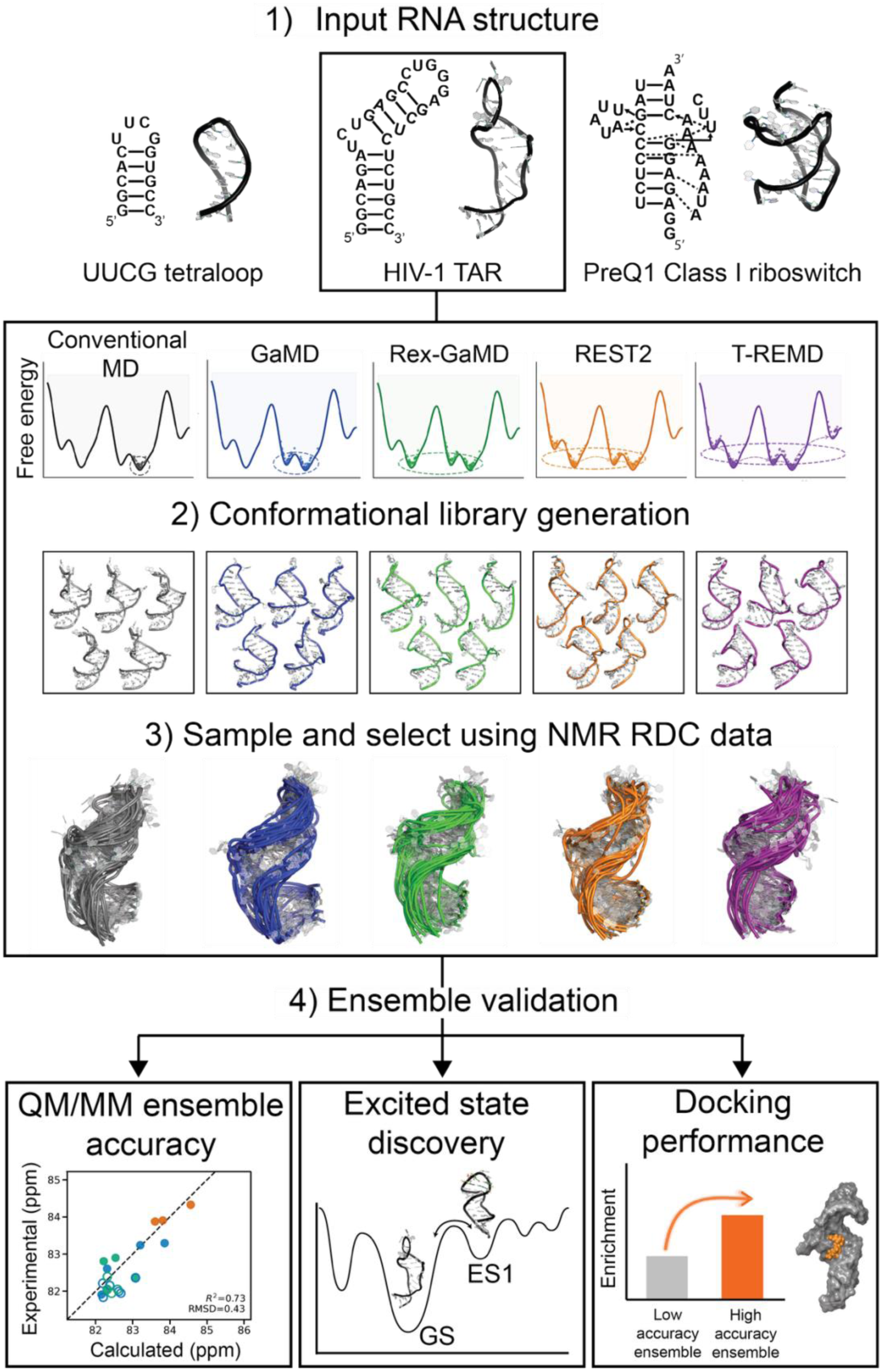
Enhanced-sampling MD workflow for atomistic RNA ensemble construction and validation. Three structurally distinct RNA systems were used in this study, including the UUCG tetraloop (PDB ID: 2KOC), HIV-1 TAR (PDB ID: 1ANR), and the preQ_1_ class I riboswitch (PDB ID: 2L1V). Starting from the input RNA structures, conformational libraries were generated using conventional MD (cMD), GaMD, Rex-GaMD, REST2, and T-REMD. The schematic free-energy profiles illustrate the expected increase in barrier crossing and conformational coverage across the sampling methods, with replica-exchange approaches enabling broader exploration of alternative RNA conformations. For each method, the generated conformational library was refined using an RDC-guided sample-and-select procedure to construct atomistic RNA ensembles. The resulting ensembles were then evaluated through independent QM/MM chemical shift validation, recovery of rare TAR conformational substates, and ensemble-based virtual screening performance. Method-specific colors are used consistently throughout the workflow, with cMD in gray, GaMD in blue, Rex-GaMD in green, REST2 in orange, and T-REMD in purple.

We show that replica-exchange methods, particularly broad-window T-REMD, produce the most accurate TAR ensembles by RDC and QM/MM chemical-shift validation, expand the accessible interhelical conformational landscape, and sample ES1-like and U23-A27-U38 base-triple conformations without predefining these states during library generation, and increase enrichment in molecular docking screens. Applying the same workflow to the preQ_1_ class I riboswitch (PDB ID: 2L1V) and the UUCG tetraloop (PDB ID: 2KOC) further supports the broader utility of enhanced-sampling-based RNA ensemble construction across structurally distinct RNAs^39,40^. The determination of more diverse and accurate ensembles enables deeper insight into how RNA substates contribute to biological function as well as high-efficiency RNA-targeted structure-based drug design.

## Results

Previous studies established that RNA 3D ensembles can be constructed by selecting conformers from computational libraries to reproduce experimental RDCs, using libraries generated by either cMD or FARFAR-NMR with QM/MM chemical-shift validation^25,30,37^. Applications to HIV-1 TAR, including ES1 modeling with an experimentally defined excited-state secondary structure, showed that ensemble accuracy strongly depends on the representativeness of the starting conformational library^14^. We therefore hypothesized that enhanced-sampling MD could provide a more representative starting library by exploring a broader and more physically continuous region of RNA conformational space than cMD, while avoiding the need to predefine an excited-state secondary structure. Because MD simulations are typically initiated from a ground-state structure, sampling alternative conformations requires crossing free-energy barriers between distinct 3D structures and base-pairing arrangements. Enhanced-sampling methods are designed to facilitate these transitions without specifying the alternative states in advance.

### Conformational libraries from enhanced-sampling MD improve RDC-refined TAR ensembles

To test this hypothesis, we generated HIV-1 TAR conformational libraries from 1000 ns simulations using cMD, GaMD, Rex-GaMD, REST2, and T-REMD, together with structural libraries generated by FARFAR2 and AlphaFold 3. Each library was subjected to the same RDC-guided ensemble optimization procedure used in previous studies^14,25,30,37^. Briefly, Prediction of ALignmEnt from Structure (PALES) was used to back-calculate RDCs for each conformer, and the sample-and-select (SAS) procedure identified subensembles whose ensemble-averaged predicted RDCs best matched the experimental measurements. We assessed ensemble accuracy by the agreement between experimental and predicted RDCs. Using the previously reported FARFAR-NMR TAR ensemble as a benchmark (Figure 2a and 2b), we found that the different libraries varied substantially in their ability to reproduce experimental RDCs. Among the methods shown in Figure 2c, T-REMD produced the strongest overall agreement with the experiment data, with an R^2^ of 0.98 and an RMSD of 1.89 Hz. REST2 and Rex-GaMD also performed well, yielding R^2^ values of 0.98 and 0.97 and RMSDs of 2.19 and 2.42 Hz, respectively. Both methods improved upon the previous FARFAR-NMR result, which gave an R^2^ of 0.96 and an RMSD of 3.14 Hz. By contrast, cMD showed performance comparable to FARFAR2, whereas GaMD was less accurate overall. The library derived by AlphaFold 3 performed substantially worse than all MD-based approaches, with an R^2^ of 0.79 and an RMSD of 7.05 Hz. Region-specific analysis further showed that T-REMD gave the lowest RDC RMSD for Helix I and the bulge, whereas REST2 gave the lowest RMSD for Helix II. Rex-GaMD also consistently improved the quality of GaMD-derived conformational libraries. These results suggest that the improved RDC agreement arises not simply from applying an enhanced-sampling label, but from increasing the breadth and representativeness of the sampled conformational pool.

**Figure 2.**
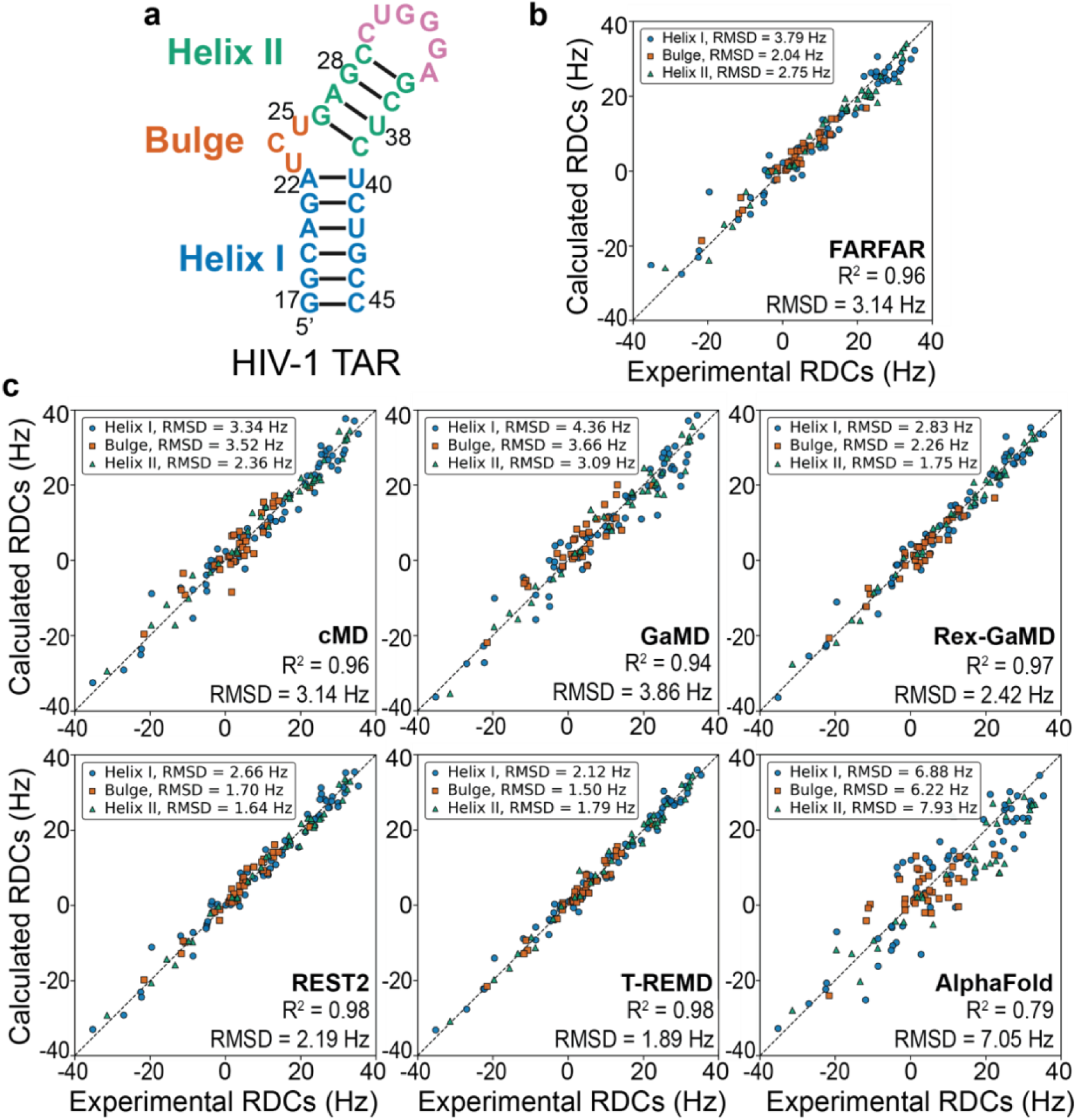
Using Enhanced-sampling MD to determine HIV-1 TAR ensemble. **(a)** Secondary structure of HIV-1 TAR. Helix I, bulge, Helix II, and loop regions are colored blue, orange, green, and pink, respectively. **(b)** Correlation between experimental and back-calculated RDCs for the previously reported FARFAR-NMR TAR ensemble^30^. **(c)** Correlations between experimental and back-calculated RDCs for ensembles generated from cMD, GaMD, Rex-GaMD, REST2, T-REMD, and AlphaFold 3 conformational libraries after the same RDC-guided sample-and-select procedure. Each point represents one RDC measurement, and colors indicate the structural region defined in panel **a**. We next examined how the extent of sampling within T-REMD influenced ensemble quality by comparing three replica temperature windows spanning 298 to 330 K, 298 to 380 K, and 298 to 480 K. The broadest temperature range is referred to as T-REMD in the main figures. Broader temperature windows allow replicas to overcome higher energetic barriers and explore wider regions of conformational space, providing a direct test of how sampling breadth affects the initial library. Consistent with this interpretation, narrowing the temperature range progressively reduced ensemble accuracy, whereas the broadest temperature range consistently gave the strongest agreement with experiment, as shown in Figure S1. These results link improved RDC agreement to the extent of conformational space sampled and identify broad-window T-REMD as the most effective strategy tested here for capturing TAR dynamics.

### Ensemble-averaged QM/MM chemical shifts independently validate the enhanced-sampling-derived TAR ensembles

We next asked whether the observed accuracy of the RDC-refined ensembles was also supported by an independent experimental observable through ensemble-averaged QM/MM chemical shift calculations. The C1′ chemical shift correlations shown in Figure 3a separated the ensembles derived from different conformational libraries. T-REMD gave the strongest overall C1′ agreement with experiment when considering both correlation and error, with an R^2^ of 0.69 and an RMSD of 0.64 ppm. REST2 gave a comparable result, with an R^2^ of 0.61 and a slightly lower RMSD of 0.58 ppm. Rex-GaMD also performed well, with an R^2^ of 0.61 and an RMSD of 0.65 ppm. FARFAR-NMR remained competitive, with an R^2^ of 0.54 and an RMSD of 0.66 ppm, whereas cMD and GaMD showed substantially weaker correlations. The ensemble derived from AlphaFold 3 showed little predictive power by this metric, with an R^2^ of 0.01 and an RMSD of 1.06 ppm. This limitation may partly reflect the far smaller number of experimentally determined RNA structures available for model training compared with protein structures in AlphaFold 3. The remaining atom-type-resolved analysis, shown in Figure S2, followed the same overall trend and further indicated that T-REMD provided the most consistently accurate description across nucleus types.

**Figure 3.**
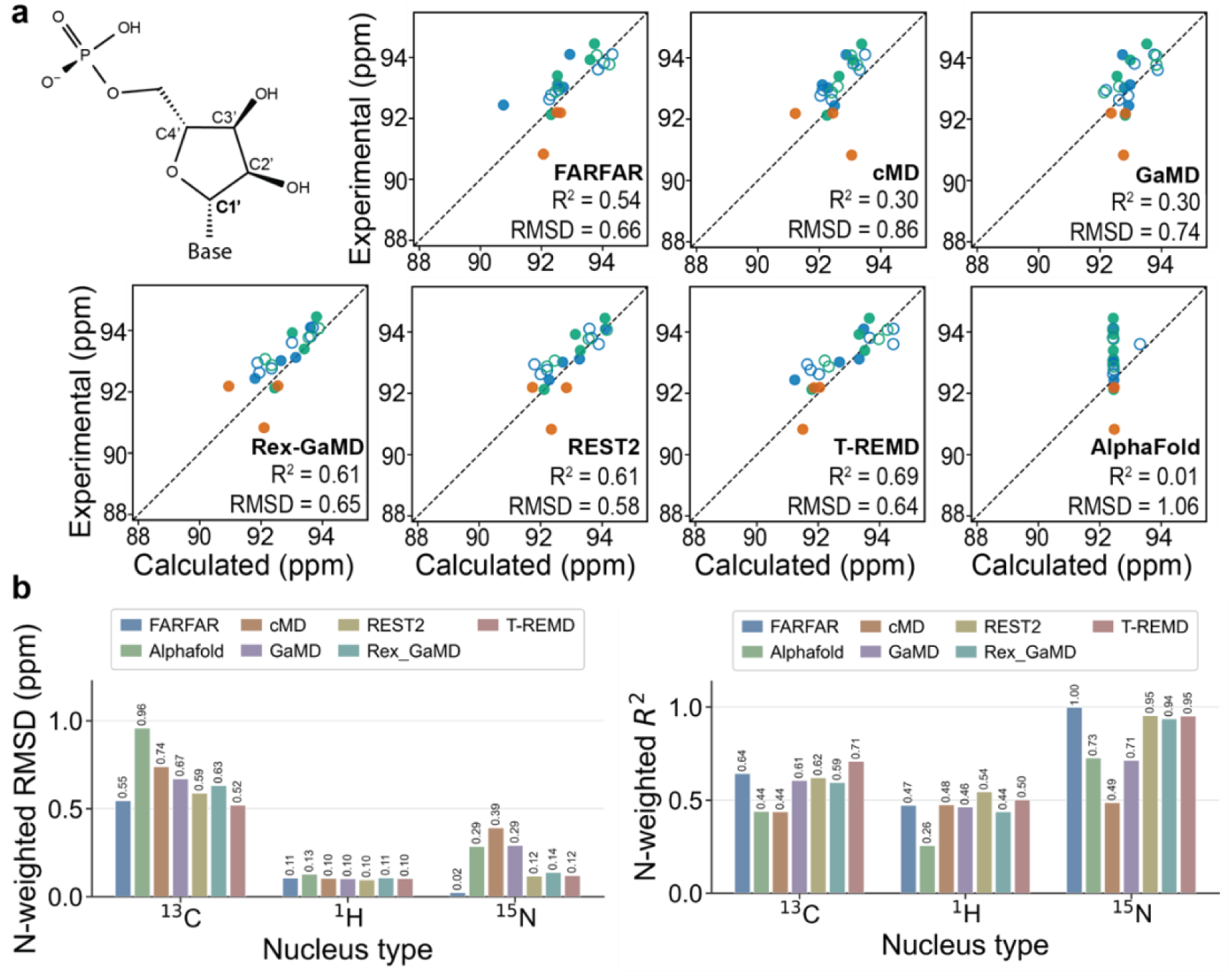
Evaluating enhanced-sampling-derived TAR ensembles using QM/MM chemical shifts. **(a)** Chemical structure of the ribose moiety with the C1′ position highlighted, together with correlations between experimental and ensemble-averaged QM/MM-predicted C1′ chemical shifts for TAR ensembles generated from FARFAR, cMD, GaMD, Rex-GaMD, REST2, T-REMD, and AlphaFold 3 conformational libraries. Each point represents one assigned C1′ chemical shift, with colors indicating TAR structural regions as defined in Figure 2a. **(b)** N-weighted RMSD and N-weighted R2 values summarizing chemical shift agreement across ^13^C, ^1^H, and ^15^N nucleus types. N-weighted metrics account for the number of assignments within each atom type, giving greater weight to more extensively measured chemical shift classes. Lower N-weighted RMSD and higher N-weighted R^2^ indicate better overall agreement with the experiment.

This ranking was further supported by the N-weighted summary metrics shown in Figure 3b, which integrate performance across the full chemical shift dataset. For ^13^C chemical shifts, which comprise the largest number of measurements and therefore provide the most statistically robust comparison, T-REMD gave the lowest N-weighted RMSD and the highest N-weighted R^2^, outperforming FARFAR and all other MD-derived ensembles. In contrast, ^1^H chemical shifts were less differentiating among methods, with N-weighted RMSDs clustered narrowly between 0.10 and 0.13 ppm, although REST2 and T-REMD still gave the highest N-weighted R^2^ values. For ^15^N, FARFAR-NMR showed the strongest agreement with experiment, but the replica-exchange ensembles also remained highly consistent with the data and clearly outperformed AlphaFold 3 and cMD. Because the ^15^N dataset contains the fewest measurements, this metric has lower resolving power than the ^13^C analysis. Taken together, the chemical shift analysis independently corroborated the RDC-based ranking and showed that the enhanced-sampling-derived ensembles, particularly those from T-REMD and REST2, provide the most accurate overall atomistic description of the TAR ensemble.

### Replica-exchange methods broaden sampling of the TAR Euler-angle landscape

To investigate the structural basis for the improved performance of the replica-exchange ensembles, we compared the global inter-helical conformational landscapes of the TAR ensembles using the three Euler angles *α*, *β*, and *γ*. In this framework, *β* reports inter-helical bending, whereas *α* and *γ* describe relative helix orientation and twist, as illustrated in Figure 4a. This Euler-angle framework was used in the earlier TAR ensemble studies to characterize global inter-helical motions^14,25,30,37^. The population maps in Figure 4b revealed clear differences in conformational coverage across methods. The AlphaFold 3 ensemble occupied a highly restricted region of *α*-*β*, *β*-*γ*, and *α*- *γ* space, indicating limited exploration of alternative global orientations. cMD and GaMD modestly expanded this coverage, and their populations remained relatively compact. In contrast, Rex-GaMD, REST2, and especially T-REMD sampled substantially broader and more continuous regions of Euler-angle space across all three projections, a trend that was also evident in the corresponding 3D ensemble representations. This broadening was most apparent along *β*, consistent with enhanced sampling of interhelical bending. It was also accompanied by broader *α* and *γ* distributions, indicating more extensive exploration of relative helix orientation and twist. The violin plots in Figure 4c recapitulated the same trend, with the narrowest angle distributions observed for AlphaFold 3 and the broadest for the replica-exchange methods. Together, these results indicate that the improved agreement with experimental RDCs and chemical shifts is associated with more complete sampling of the global TAR conformational landscape.

**Figure 4.**
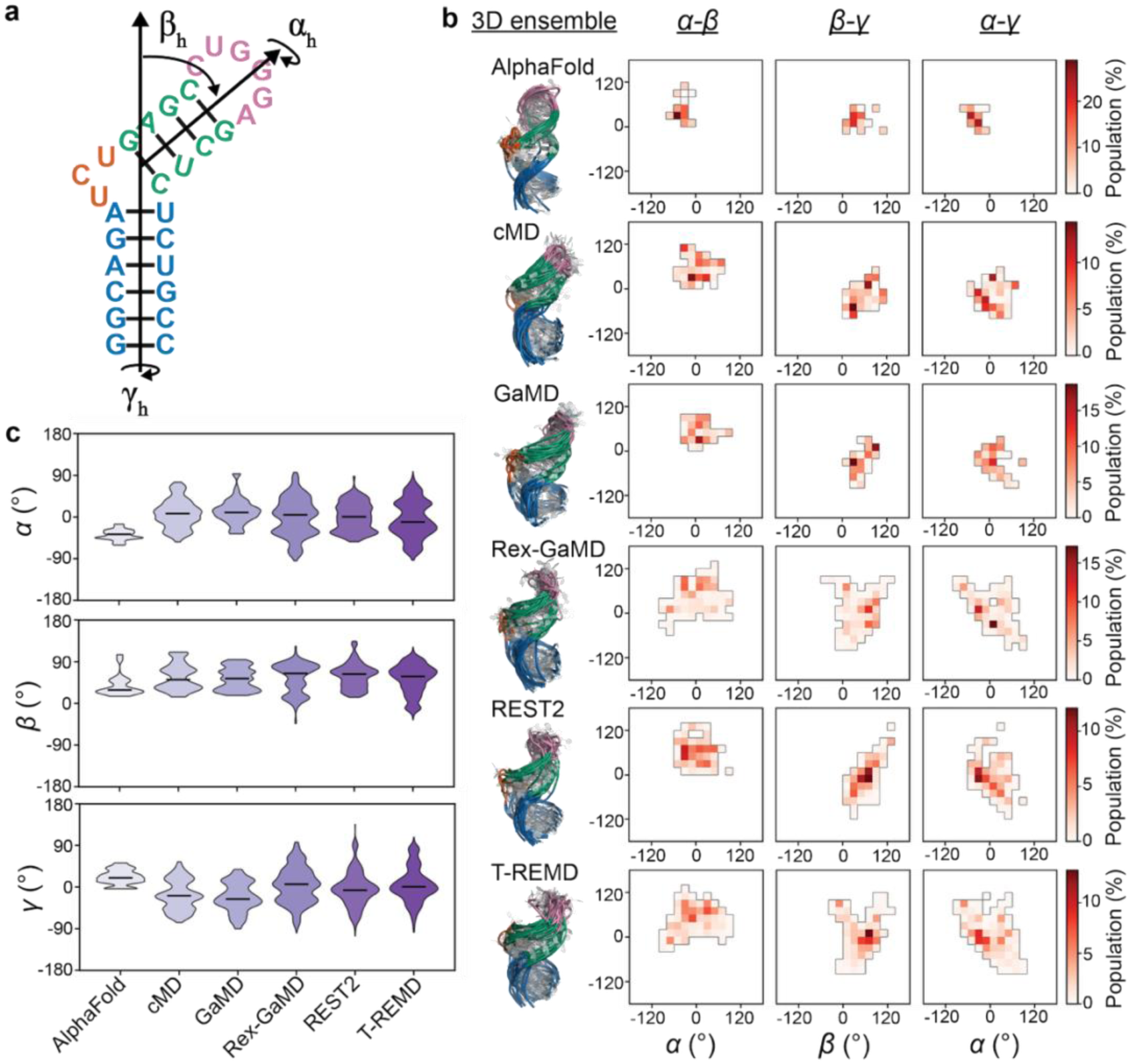
Replica-exchange methods broaden the interhelical conformational landscape of HIV-1 TAR. **(a)** Schematic of the TAR secondary structure and the Euler angles used to describe global motions between Helix I and Helix II. **(b)** 2D population maps and 3D ensemble overlays showing the conformational coverage of each method. **(c)** Violin plots showing the distributions of α, β, and γ for ensembles generated by AlphaFold 3, cMD, GaMD, Rex-GaMD, REST2, and T-REMD.

### Broad replica-exchange sampling directly captures low-populated excited states of TAR

Prior NMR relaxation-dispersion studies identified two well-characterized TAR excited states, including ES1, an apical-loop rearrangement populated at approximately 13%, and ES2, a much rarer state populated at approximately 0.4% that remodels the bulge, upper stem, and apical loop under the reported solution conditions^11,13,36^. Productive Tat recognition also requires formation of the U23-A27-U38 base triple. We next asked whether the broader conformational coverage achieved by the enhanced-sampling MD libraries was sufficient to sample conformations resembling low-populated TAR substates without imposing these states during conformational library generation. The outcome depended strongly on the sampling protocol. The secondary structures corresponding to the ground state, ES1, and the U23-A27-U38 base-triple state are shown in Figure 5a. Neither cMD nor GaMD sampled ES1 or the base-triple state, whereas Rex-GaMD sampled the base triple with an observed frequency of 0.89%, and REST2 sampled ES1 with a frequency of 0.39% (Figure 5b). T-REMD provided greater access to both rare states. ES1 was observed at frequencies of 3.44%, 2.00%, and 5.62% for the 298 to 330 K, 298 to 380 K, and 298 to 480 K conditions, respectively. The variation among these values indicates that the 1 μs simulations were insufficient to determine a converged equilibrium population. In comparison, base-triple sampling increased from undetected in the 298 to 330 K condition to 3.37% and 3.87% in the 298 to 380 K and 298 to 480 K conditions, respectively, consistent with broader temperature ranges facilitating conformational barrier crossing. The goal of this analysis was to evaluate whether each sampling method could access these rare TAR substates rather than to quantify their equilibrium populations. Accordingly, the reported frequencies describe state occurrence within the finite trajectories and should not be interpreted as equilibrium populations. Representative T-REMD structures exhibited the expected ES1 and base-triple topologies (Figure 5c). We note that the experimentally characterized ES1 is stabilized by a protonation-dependent C30-A+35 wobble involving A35, together with a U31-G34syn pair^11^. Because our simulations used a fixed-protonation force field with neutral A35, the identified conformers cannot fully reproduce the experimental ES1 and are therefore best regarded as ES1-like structural mimics.

**Figure 5.**
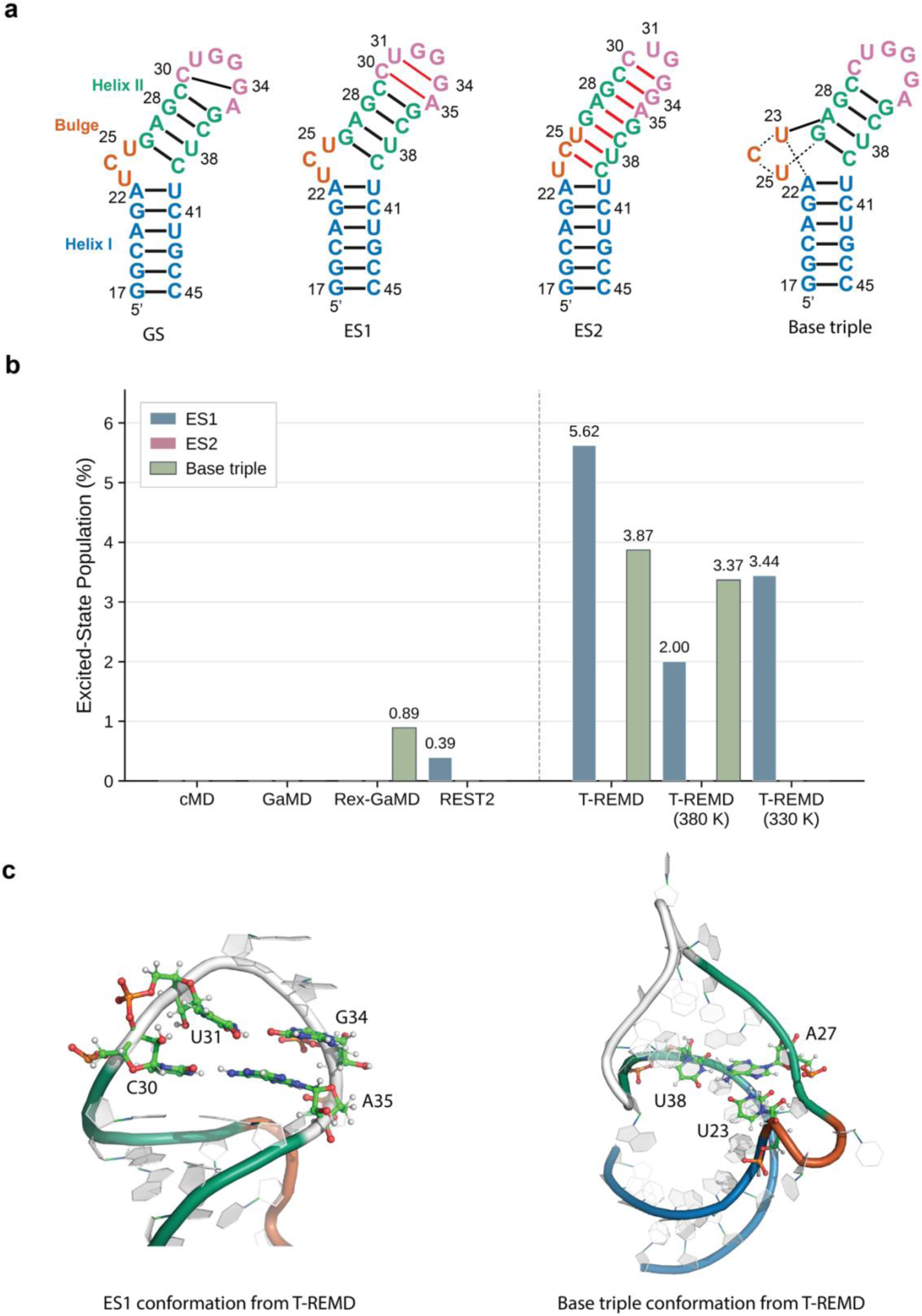
Broad T-REMD directly recovers low-populated TAR conformational substates. **(a)** Secondary structures of the TAR ground state, ES1, ES2, and U23-A27-U38 base-triple state. Black bars represent canonical Watson-Crick-Franklin base pairs in the GS, red bars represent new base pairs formed in the alternative conformations. **(b)** Populations of ES1, ES2, and base-triple conformations detected from different conformational libraries. **(c)** Representative ES1 and base-triple conformations recovered directly from the T-REMD ensemble.

Unlike prior TAR excited-state ensemble studies, in which an experimentally inferred ES1 secondary structure was supplied during FARFAR library generation, the ES1-like and base-triple conformations identified here emerged directly from MD sampling without pre-specification. We also searched the complete trajectories before SAS for the experimentally characterized ES2 base-pairing pattern but did not detect ES2-like conformers under the present simulation conditions. Expanding the T-REMD temperature range may facilitate crossing of larger conformational barriers and increase the likelihood of sampling ES2-like structures, although this possibility remains to be tested. Even if ES2 conformers are generated in the raw library, however, their very low experimental population of approximately 0.4% produces little measurable contribution to ensemble-averaged RDCs, making them difficult to recover through RDC-guided ensemble refinement alone^13^.

### More accurate TAR ensembles better capture ligand-bound conformations and improve ensemble-based virtual screening performance

To assess the functional significance of ensemble accuracy, we asked whether the simulated TAR ensembles captured experimentally observed ligand-bound interhelical orientations and whether this was accompanied by improved ensemble-based virtual screening performance. The experimental reference set comprised six TAR-small-molecule complex structures, 1ARJ, 1LVJ, 1QD3, 1UTS, 1UUD, and 1UUI, which span diverse ligand-bound orientations, as shown in Figure 6a. We first evaluated the compatibility of each ensemble with these structures using Mahalanobis distance in Euler-angle space^41^. Rex-GaMD and T-REMD gave the lowest mean Mahalanobis distances across the six complexes, with values of 1.11 and 1.19, respectively, compared with 1.48 for FARFAR-NMR and 1.55 for the 1ANR NMR ensemble (Figure 6b). Because Mahalanobis distance is normalized by ensemble covariance and is therefore influenced by ensemble breadth, we also calculated the minimum geodesic rotation distance between each complex and any ensemble conformer, together with the fraction of conformers within a 20° cutoff. FARFAR-NMR and Rex-GaMD gave comparable mean nearest distances of 12.11° and 12.70°, respectively, while Rex-GaMD provided the most uniform coverage, with at least one conformer within 20° of all six complexes. GaMD and REST2 also showed strong direct coverage, with mean nearest distances of 12.88° and 14.57°, respectively, and at least one conformer within 20° of five of the six complexes. T-REMD reduced the mean nearest distance from 20.22° for cMD to 15.64° and covered four of the six complexes within 20°. Together, these analyses indicate that the low Mahalanobis distances of T-REMD and Rex-GaMD reflect their broader Euler-angle distributions, while the direct distance metrics demonstrate that enhanced-sampling ensembles generally improve access to experimentally observed ligand-bound interhelical orientations relative to cMD.

**Figure 6.**
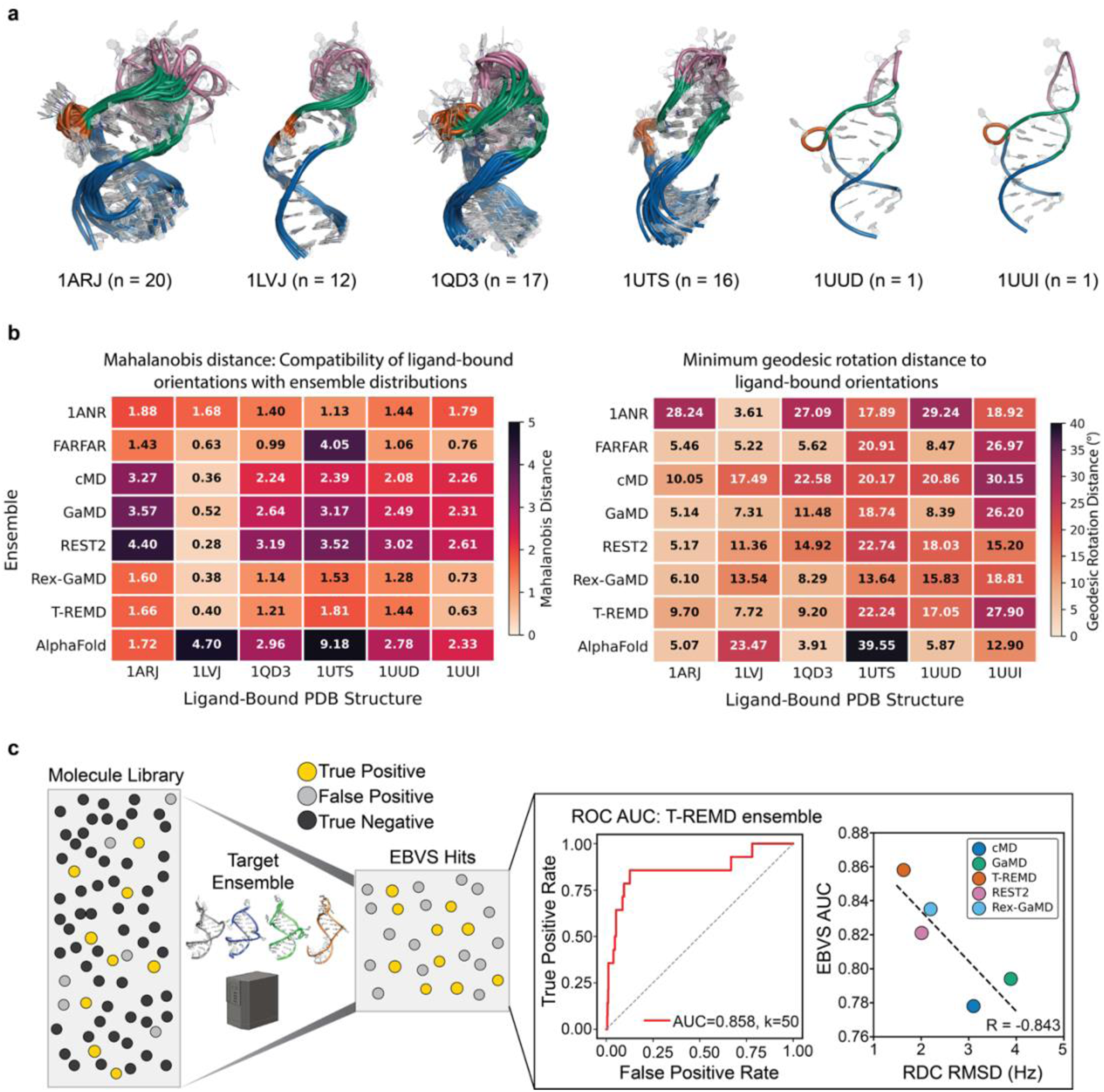
Experimental ligand-bound TAR coverage and functional evaluation of enhanced-sampling ensembles. **(a)** Experimental ligand-bound HIV-1 TAR structures. **(b)** Heatmaps comparing the TAR ensembles with experimentally observed ligand-bound interhelical orientations in Euler-angle space. The left heatmap shows the Mahalanobis distance, with lower values indicating greater compatibility with the center and breadth of the corresponding ensemble distribution. The right heatmap shows the minimum geodesic rotation distance between any conformer in each ensemble and any experimental model in the corresponding ligand-bound PDB structure, with lower values indicating closer direct access to the experimental interhelical orientation. **(c)** Functional evaluation of the RNA ensembles by EBVS. The left panel shows the ROC AUC curve obtained using the ensemble from T-REMD. The right panel shows the correlation analysis between RDC RMSD and EBVS AUC across MD-derived ensembles.

The practical value of recovering ligand-bound-like conformations depends on whether docking against these structures can distinguish known TAR binders from non-hits. We next tested whether improved structural agreement translated into improved ensemble-based virtual screening (EBVS) performance. Previous work curated a library of small molecules that have been experimentally tested for their ability to bind TAR^42^. This library includes 14 known hits and 637 known non-hits. Figure 6c shows a Receiver Operating Characteristic (ROC) curve for the T-REMD-derived ensemble, which achieved an area under the curve (AUC) of 0.86. The corresponding curves for the remaining ensembles are shown in Figure S3. Across the MD-derived ensembles, EBVS performance was broadly consistent with ensemble accuracy. T-REMD, which yielded the lowest RDC RMSD, also produced the highest EBVS AUC of 0.86. REST2 and Rex-GaMD showed intermediate behavior, with AUC values of 0.82 and 0.83 at RDC RMSDs of approximately 2.0 to 2.2 Hz. In contrast, cMD and GaMD, which had larger RDC RMSDs, gave lower AUC values of 0.78 and 0.79, respectively.

Across the five MD-derived ensembles, EBVS AUC showed a strong inverse trend with RDC RMSD, with an R value of -0.843. This trend is consistent with the possibility that ensembles that more accurately reproduce experimental RDCs may also provide improved enrichment of known TAR binders, although the limited number of methods precludes a statistically robust conclusion. Together, these results suggest that improved TAR ensemble accuracy may have functional consequences, including better coverage of experimentally observed ligand-bound conformations and improved virtual screening performance.

### Enhanced-sampling ensemble determination generalizes to structurally distinct RNAs

Finally, we investigated whether the advantages of enhanced-sampling-based ensemble determination that we have observed generalize beyond TAR to structurally distinct RNA systems. We applied the same workflow to two additional RNAs with different folds and dynamical features, the preQ_1_ class I riboswitch with PDB ID 2L1V and the UUCG tetraloop with PDB ID 2KOC^39,40^. For both systems we benchmarked the methods for ensemble accuracy and ability to sample alternative conformations starting from the ground state as with TAR. We were unable to perform the EBVS benchmark as there was not a suitable small molecule dataset for these RNAs. For the preQ_1_ class I riboswitch, cMD produced an ensemble that was broadly consistent with the experimental RDCs, with an overall R^2^ of 0.952 and an RMSD of 2.170 Hz, as shown in Figure 7a. T-REMD further improved the agreement to R^2^ of 0.970 with an RMSD of 1.704 Hz, indicating a more accurate ensemble. The corresponding 3D ensembles further support this conclusion. Whereas the cMD ensemble remains relatively compact and predominantly captures the stacked state, the T-REMD ensemble samples a broader conformational landscape and includes an unstacked conformation consistent with the experimentally observed state, as shown in Figure 7b.

**Figure 7.**
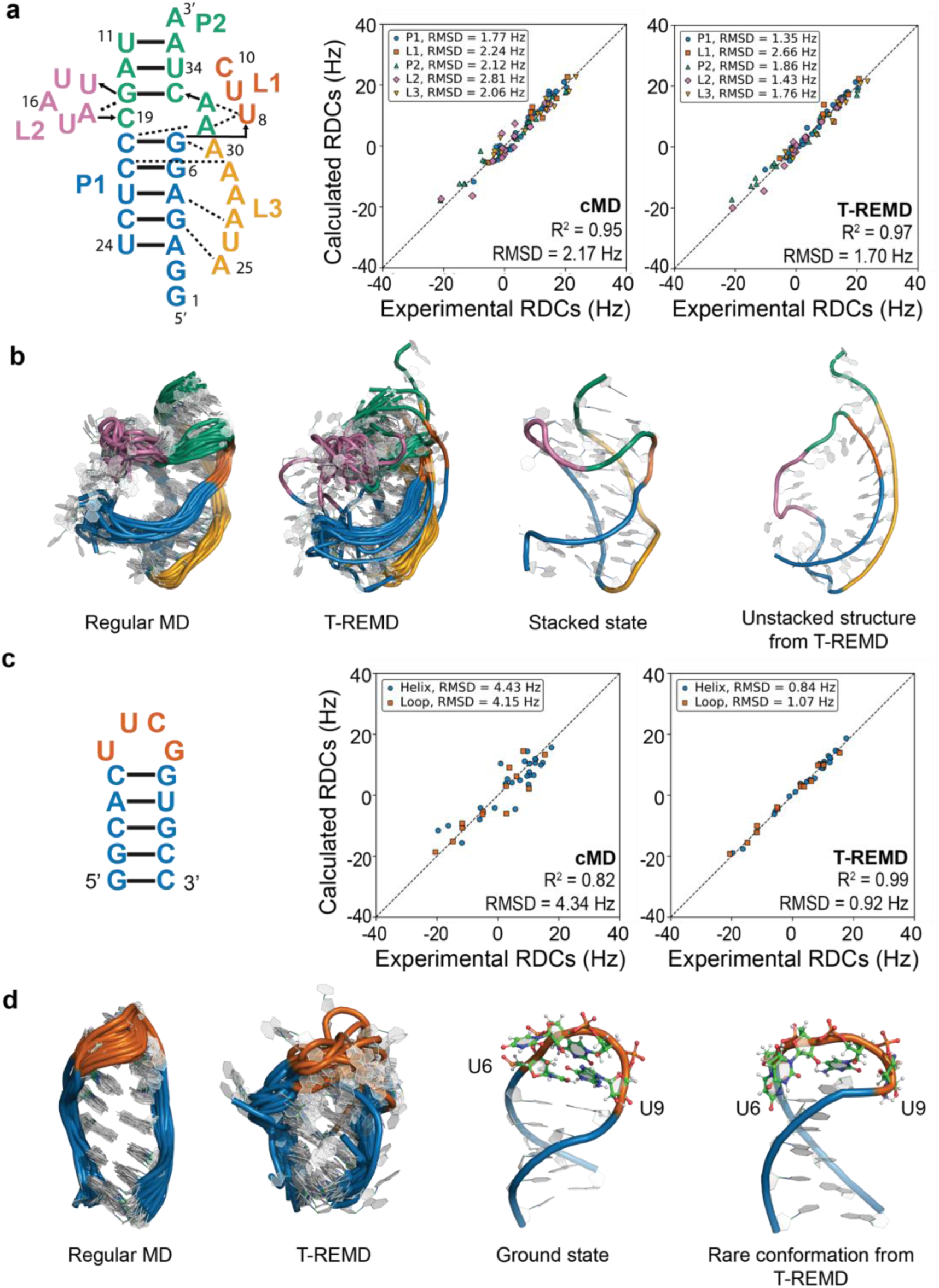
Enhanced-sampling MD simulations improve RNA ensemble accuracy across two additional, structurally distinct RNAs. **(a)** Secondary structure of the preQ_1_ class I riboswitch and RDC correlation plots for ensembles generated from cMD and T-REMD. Points are colored by structural region. **(b)** 3D ensembles of preQ_1_ class I riboswitch generated from cMD and T-REMD, along with representative stacked and unstacked conformations. **(c)** Secondary structure of the UUCG tetraloop and RDC correlation plots for cMD and T-REMD ensembles. Points are colored by helix and loop regions. **(d)** 3D ensembles of UUCG tetraloop generated from cMD and T-REMD, along with representative ground-state and rare loop conformations.

The remaining enhanced-sampling methods showed mixed but generally improved performance relative to cMD. As shown in Figure S4, GaMD gave weaker overall agreement than cMD, with an R^2^ of 0.914 and an RMSD of 2.912 Hz. Rex-GaMD and REST2 both improved the agreement, yielding R^2^ values of 0.960 and 0.961 and RMSD values of 1.971 and 1.942 Hz, respectively. The corresponding ensemble representations are also shown in Figure S4. Together, these results indicate that broader conformational sampling improves the preQ_1_ riboswitch ensemble, with T-REMD providing the strongest overall performance among the methods tested.

A similar trend was observed for the UUCG tetraloop. cMD showed only moderate agreement with the experimental RDCs, with an overall R^2^ of 0.825 and an RMSD of 4.341 Hz, as shown in Figure 7c. In contrast, T-REMD yielded a much more accurate ensemble, improving the agreement to an R^2^ of 0.992 and an RMSD of 0.923 Hz. The corresponding 3D ensembles again revealed a clear difference between the two methods. Whereas the cMD ensemble remains close to a dominant loop geometry and primarily captures the ground-state conformation, the T-REMD ensemble accesses a rare loop conformation consistent with the experimentally observed alternative state, as shown in Figure 7d. These results indicate that T-REMD captures both the RDC-defined ensemble and the experimentally relevant conformational heterogeneity of the UUCG tetraloop more effectively than cMD.

The remaining enhanced-sampling methods again improved upon cMD but did not match the overall performance of T-REMD. As shown in Figure S5, GaMD and Rex-GaMD substantially improved the UUCG RDC agreement relative to cMD, with R^2^ values of 0.976 and 0.981 and RMSD values of 1.642 and 1.450 Hz, respectively. REST2 gave a more moderate improvement, with an R^2^ of 0.930 and an RMSD of 2.729 Hz. The corresponding conformational ensembles are shown in Figure S5. Taken together, these results show that the advantage of T-REMD is not limited to TAR but extends to structurally distinct RNA systems with different folds and dynamical modes.

## Discussion

A central conclusion of this study is that the quality of an RNA ensemble is determined first by the quality of the starting conformational library and only then by the experimental refinement used to select or reweight conformers. For highly dynamic RNAs such as TAR, experimental observables report on averages over many conformational states. Refinement therefore cannot recover structurally relevant states that are absent from the initial conformational pool. This observation is consistent with the broader shift in RNA structural biology from static structures toward ensemble-based descriptions of folding, recognition, and function, including the growing recognition that low-populated excited conformational states can have important _functional roles9,10,12–14,30,43._

Our results show that the accuracy of MD-derived RNA ensembles is not determined by the force field alone. It depends on the full simulation model, including the force field, water model, and the extent and character of conformational sampling. Earlier TAR ensemble studies showed that broader conformational libraries and longer simulations can improve agreement with RDCs, whereas incomplete sampling limits the utility of subsequent ensemble selection^25,38^. In the present study, all simulations were performed using the four-point OPC water model. Compared with prior studies using the same RNA force field but a less optimized water model, our cMD simulations already showed improved agreement with experiment. This suggests that the observed performance reflects not only the enhanced-sampling protocol, but also the underlying physical model used to describe the molecular interactions. Accordingly, the performance of each method should be interpreted as the combined outcome of its sampling strategy, force field, and solvent model, rather than as a property of the force field alone. More broadly, prior work has shown that water-model choice can strongly influence RNA conformational behavior, underscoring the importance of selecting an appropriate force-field and solvent-model combination^27,29,44^.

Enhanced-sampling MD and FARFAR provide complementary routes to RNA conformational sampling. FARFAR rapidly generates conformers consistent with a specified secondary structure, whereas enhanced-sampling MD allows base-pairing rearrangements to emerge directly from the trajectories while preserving the energetic coupling between local and global motions in explicit solvent and ions. This distinction is particularly relevant for TAR, where bulge rearrangements are coupled to large-scale interhelical reorientation. For very lowly-populated states, however, specifying the excited-state secondary structure may still be necessary because their limited contribution to ensemble-averaged RDCs is insufficient to guide refinement. Thus, our results complement prior FARFAR-NMR studies by showing that some rare TAR substates can emerge without prior structural specification.

A further limitation applies to both FARFAR and AlphaFold 3. FARFAR relies on fragment-based structural priors and secondary-structure constraints, whereas AlphaFold 3 cannot predict structures of RNAs containing non-standard nucleotides; consequently, neither method is inherently suited to determining how chemical modifications reshape RNA conformational landscapes. This limitation is important because even subtle modifications can alter local sugar-pucker, stacking, hydrogen-bonding, and kinetic preferences, thereby redistributing the global ensemble. Previous work showed that 2′-O-methylation at TAR bulge residues shifts the conformational equilibrium in a measurable and cooperative manner, and broader studies of methylation support the idea that small chemical changes can reshape nucleic acid conformational landscapes^45^. Enhanced-sampling MD is therefore likely to be especially useful for modified RNAs, where local chemical changes must propagate through a continuous and environment-dependent energy landscape. The same consideration may extend to RNAs containing specialized structural features, such as the 5′ cap guanosine in the HIV-1 5′ UTR, whose functional structural consequences may be difficult to represent using standard fragment libraries or secondary-structure-guided modeling alone^46^.

An important test of library quality is whether a method generates conformations relevant to excited states. Recent work on TAR has shown that excited states are not single structures, but subensembles with characteristic secondary-structure features, internal heterogeneity, and experimentally measurable populations^9,12–14,37,47^. Capturing such states therefore requires more than identifying one rare frame. It requires recovering the conformational family from which the state is built. In this context, the ability of the best-performing methods in our study to recover ES1-like and base-triple-related conformations is particularly important. These states provide a demanding test of whether the simulation library contains the structurally and functionally relevant minority substates needed for realistic ensemble construction.

The relationship between ensemble accuracy and EBVS performance should also be viewed as a connected problem. TAR has long served as a model for adaptive recognition, and prior work showed that free TAR can dynamically access conformations resembling ligand- or protein-bound states, while alternative excited conformations can sequester recognition elements and alter binding competence^38,42^. From that perspective, the link between ensemble accuracy and downstream screening performance is not incidental. A more accurate ensemble should provide a more realistic distribution of binding-competent and binding-incompetent conformers, thereby improving the physical basis for ensemble-based virtual screening in RNA-targeted drug discovery. At the same time, EBVS performance depends on several methodological factors beyond ensemble quality, including the docking program, scoring function, ensemble size, receptor preparation, ligand preparation, and scoring protocol. The EBVS results presented here should therefore be interpreted as evidence that improved RNA ensembles can enhance downstream screening, rather than as a complete benchmark of all determinants of docking performance. Those parameters are more appropriately addressed in a dedicated EBVS benchmarking study.

Importantly, even the best-performing method in this comparison does not define a complete TAR free-energy landscape. Our benchmarks identify T-REMD as the most useful starting conformational library among the methods tested, but they do not validate the accuracy or biological relevance of the entire sampled landscape. Moreover, sampling a conformation does not by itself establish that it is a genuine excited state. ES1-like conformations could be recognized because ES1 was already known. Without this prior knowledge, they might not have been distinguishable from the many other structures generated. This limits the current framework’s capacity for *de novo* excited-state discovery and highlights the need for additional experimental or energetic criteria to identify physically relevant rare states. This challenge will likely increase with RNA size and tertiary complexity, especially for systems involving long-range contacts and highly cooperative tertiary interactions. Although broader T-REMD temperature windows can expand sampling, this improvement comes with substantial computational cost (Figure S6). Future work will therefore require strategies that preserve the physical advantages of enhanced sampling while reducing the number of replicas or improving the efficiency of rare-state discovery.

In summary, our results support a view of RNA ensemble determination in which simulation and experiment play complementary roles. Simulation is most valuable when it generates a rich and physically plausible conformational library, whereas experiment is essential for identifying which parts of that pool correspond to the true solution ensemble. For TAR, this framework highlights the importance of the solvent model, sampling strategy, local bulge and loop rearrangements, and the ensemble nature of excited states. More broadly, it suggests that future progress in RNA structural biology will depend on integrating force field, water model, and enhanced sampling into a unified workflow for determining RNA conformational landscapes at atomic resolution.

## Conclusion

This work demonstrates that RNA ensemble determination is strongly shaped by the quality of the initial conformational library and that enhanced-sampling MD can improve this step relative to cMD and structure-prediction-based libraries. For HIV-1 TAR, T-REMD generated the most accurate RDC-refined ensemble among the methods tested and was independently supported by ensemble-averaged QM/MM chemical shifts. REST2 and Rex-GaMD also improved ensemble quality relative to cMD, indicating that replica-exchange strategies provide a more effective route to conformational library generation. The improved performance of these methods was associated with broader and more continuous sampling of the TAR interhelical conformational landscape, consistent with more complete recovery of the global motions required for accurate ensemble determination. A central advance of this study is that broad enhanced sampling, particularly T-REMD, directly recovered low-populated TAR substates, including ES1 and the U23-A27-U38 base-triple state, without requiring excited-state secondary structures to be specified during library generation. These improvements were also functionally meaningful, leading to improved coverage of experimentally observed ligand-bound TAR conformations and improved ensemble-based virtual screening performance. The same overall trend extended to the preQ_1_ class I riboswitch and UUCG tetraloop, supporting the broader applicability of this strategy to structurally distinct RNAs. Together, these results establish replica-exchange enhanced sampling, especially T-REMD, as an effective approach for constructing experimentally validated RNA ensembles and revealing low-populated functional substates directly from atomistic simulations.

## Methods

### Definitions of RNA Conformational Ensembles and States

Throughout this study, we use the term “conformational ensemble” to represent the population-weighted distribution of RNA conformations, whereas the corresponding free-energy landscape describes their relative free energies and the barriers separating conformational states. We use “conformational state” to refer to a subensemble of 3D conformations that share the same secondary structure, occupying energetic minima on this landscape. The dominant, lowest free-energy state is termed the “ground state” (GS). We use “alternative conformational state” as a general term for any structurally distinct state relative to the GS and use “excited state” (ES) for a transient, low-population alternative state that occupies a higher-free-energy basin and exchanges with the GS. Alternative states including ESs may differ from the GS in secondary structure, three-dimensional structure, or both.

### Overview of Enhanced-Sampling Methods

Replica-exchange methods run multiple replicas in parallel and periodically attempt exchanges between neighbors under an acceptance criterion that preserves the corresponding equilibrium distributions, allowing configurations to visit more strongly tempered conditions, cross barriers, and return to the target condition. In temperature replica-exchange MD (T-REMD), the replicas are simulated at different temperatures, such that the entire solvated system is tempered^48^. In replica exchange with solute tempering (REST2), solvent-solvent interactions remain unscaled, whereas interactions involving the RNA are scaled to selectively temper the solute^49^. Gaussian accelerated MD (GaMD) uses a different strategy, adding a smooth harmonic boost potential when the system potential energy falls below a defined threshold and thereby reducing effective barriers without requiring predefined collective variables^50^. Replica-exchange GaMD (Rex-GaMD) combines these strategies by coupling replica exchange with replicas characterized by different GaMD boost-potential parameters^51^.

### RNA systems and structural preparation

Three RNA systems were investigated in this study, HIV-1 TAR with PDB ID 1ANR, the preQ_1_ class I riboswitch with PDB ID 2L1V, and the UUCG tetraloop with PDB ID 2KOC^39,40,52^. System preparation for MD simulation was carried out using CHARMM-GUI^53–56^. RNA molecules were described using the OL3 RNA force field and solvated with the OPC water model^31,57^. Each system was placed in an explicit solvent box extending at least 10 Å from the solute. The systems were neutralized using KCl, and the ionic strength was adjusted by adding 0.15 M KCl and 0.01 M MgCl_2_.

### Details for different molecular dynamics simulations

All MD simulations were performed using the GPU-accelerated pmemd.cuda implementation in Amber24^58^. Nonbonded interactions were evaluated with a 10 Å cutoff, and long-range electrostatic interactions were treated using the particle mesh Ewald method. All bonds involving hydrogen atoms were constrained with the SHAKE algorithm, which enabled the use of a 2 fs integration time step^59^. All simulations were initiated using the same equilibration procedure. First, a two-stage minimization was carried out. In the initial stage, harmonic restraints of 50 kcal·mol^-1^·Å^-2^ were applied to the full RNA structure, followed by an unrestrained minimization of the entire system. After minimization, the systems were heated from 0 to 298 K over 100,000 steps in the NVT ensemble while maintaining the same harmonic restraints on the RNA. This was followed by a density-equilibration stage consisting of five NPT runs with a combined duration of 0.2 ns at 298 K and 1 bar. Temperature control was achieved using Langevin dynamics with a collision frequency of 1.0 ps^-1^, and pressure was regulated using the Monte Carlo barostat^60–62^. Production simulations were then performed in the NPT ensemble for at least 1000 ns for each replica and each method. Trajectory coordinates were saved every 0.02 ns.

cMD simulations were carried out for each RNA system using the equilibration procedure described above, followed by 1000 ns of production dynamics at 298 K. The resulting trajectories were used as the conformational library for the cMD-based ensemble analysis.

GaMD simulations were performed on the equilibrated RNA systems using the GaMD module implemented in the GPU version of Amber24^63^. For each system, the GaMD protocol included a preparatory stage for collecting potential statistics to define the acceleration parameters, followed by an equilibration stage after application of the boost potential, and then production simulations of at least 1000 ns. All GaMD simulations were performed at the dual-boost level, in which one boost potential was applied to the total potential energy and the other to the dihedral potential energy. The averaging window used to collect the potential energy statistics was 240,000 steps, and the upper limits of the boost potential standard deviations were set to 3 kcal/mol for both the total and dihedral energetic terms. Trajectory coordinates were saved every 0.02 ns.

Replica-exchange Gaussian accelerated MD (Rex-GaMD) simulations were performed for all three RNA systems using 8 replicas at 298 K, following the protocol of Hasse and Huang^64^. The replicas differed in the value of the total-potential boost parameter, which ranged from 2.0 to 5.5 in increments of 0.5. Each Rex-GaMD simulation was run for at least 1000 ns. Populations reported for GaMD and Rex-GaMD were computed as raw frame frequencies on the boosted potential and were not reweighted to the canonical ensemble; they therefore indicate conformational accessibility rather than equilibrium population.

Replica exchange with solute tempering 2 (REST2) simulations were carried out in Amber24 using the REAF module^58^. For HIV-1 TAR and the preQ_1_ class I riboswitch, 32 replicas were distributed over the temperature range 298-480 K. For the UUCG tetraloop, 16 replicas were used over the same temperature range. The exchange rates were approximately 25% for all REST2 simulations. Each simulation was run for at least 1000 ns. The 298 K trajectories were extracted and used as conformational libraries for downstream ensemble construction.

For TAR, three T-REMD simulations were used to evaluate the effect of sampling breadth on ensemble quality. These simulations used temperature windows of 298 to 480 K with 80 replicas, 298 to 380 K with 40 replicas, and 298 to 330 K with 16 replicas. All three TAR T-REMD simulations have exchange rates of approximately 23%. For the preQ_1_ class I riboswitch, T-REMD was performed over 298-480 K using 64 replicas with an exchange rate of approximately 23%. For the UUCG tetraloop, T-REMD was performed over 298-480 K using 80 replicas with a similar exchange rate. Each T-REMD simulation was run for at least 1000 ns. The 298 K trajectories were extracted and used as conformational libraries for downstream ensemble construction.

An additional conformational library was generated using AlphaFold 3^24^. To increase conformational diversity, multiple independent AlphaFold 3 predictions were performed using different random seeds. In total, 50,000 conformers were generated and pooled to form the final AlphaFold 3 conformational library. This library was then analyzed using the same downstream framework applied to the MD-derived libraries.

### Residual dipolar coupling analysis

RNA ensembles were refined against experimentally measured one-bond RDCs. For HIV-1 TAR, four RDC data sets were used, corresponding to the E0, EI22, EII22, and EI3 constructs^25^. These variably elongated constructs modulate molecular alignment and provide semi-independent orientational restraints for ensemble refinement. For the preQ_1_ class I riboswitch and the UUCG tetraloop, RDC data were taken from the studies of Zhang et al. and Borkar et al., respectively^65,66^. All RDC data were analyzed within the same refinement framework.

RDCs were back-calculated using PALES with a steric alignment model^67^. For the elongated TAR constructs, conformers from the non-elongated simulation libraries were elongated in silico using idealized A-form geometry prior to PALES analysis, following earlier RNA ensemble studies^30^. The calculations were carried out assuming an effective Pf1 concentration of 0.022 g/mL within a rod-like liquid-crystalline alignment model. Predicted RDCs were obtained separately for each construct. For each ensemble, the predicted RDCs were averaged over all conformers. Construct-specific scaling factors were then applied to account for differences in overall alignment magnitude, arising in part from differences in Pf1 phage concentration among the experiments, as described previously^14,30^.

### Sample and select

Ensembles were generated from the simulation-derived conformational libraries using the sample-and-select (SAS) procedure previously described for RNA RDC ensemble construction^30,68^. A simulated-annealing Monte Carlo protocol was used to identify subsets of conformers that minimized the discrepancy between measured and predicted RDCs, with the target function defined by the squared deviation between experimental and back-calculated RDCs and including construct-specific alignment scaling factors. The initial effective temperature for simulated annealing was set to 100 and decreased by a factor of 0.9 at each step, using the same parameters and settings as reported by Shi et al. A series of SAS runs was performed over a range of ensemble sizes, starting from ensemble size of 1 and increasing until the RDC RMSD reached a plateau.

### Interhelical Euler angle calculation

Global interhelical motions were quantified using the three interhelical Euler angles *α*, *β*, and *γ*, which describe the relative orientation of two A-form helical segments^69^. In this description, *β* reports the interhelical bend angle, whereas *α* and *γ* describe the relative twist and orientation about the two helices. For each conformer, the upper and lower helices were defined using the corresponding stem base pairs and aligned to idealized A-form helical reference frames, after which the relative orientation was expressed in terms of the three Euler angles.

For TAR, helix I was defined using residues C19-G21 and helix II using residues G26-G28, consistent with previous analyses. The resulting *α*-*β*, *β*-*γ*, and *α*- *γ* distributions were used to compare global conformational coverage across methods. For comparison with experimentally determined TAR-small-molecule complexes, Euler angles were calculated for the ligand-bound structures using the same definition, and Mahalanobis distance was used to quantify the extent to which each ensemble covered the experimentally observed ligand-bound conformational space. The base-pairing mode of TAR excited states were detected by X3DNA-DSSR^70^.

### QM/MM chemical shift calculation

Ensemble-averaged NMR chemical shifts were calculated to independently validate the refined RNA ensembles using the automated fragmentation quantum mechanics/molecular mechanics (AF-QM/MM) approach^71^. Chemical shifts were predicted for each conformer individually and then averaged over all conformers in a given ensemble. The resulting ensemble-averaged values were compared directly with experiment. Following earlier RNA ensemble studies, resonance-type-specific linear corrections were applied to the predicted ensemble-averaged chemical shifts prior to comparison with measured values^14,30^.

For AF-QM/MM chemical shift calculations, each RNA conformer was first subjected to five steps of conjugate-gradient minimization with harmonic restraints of 2 kcal·mol^-1^·Å^-2^ on all heavy atoms to regularize bond lengths and angles and reduce noise in the subsequent quantum calculations. Each structure was then partitioned into quantum fragments centered on individual nucleotides and containing 2-6 neighboring nucleotides within a heavy-atom distance cutoff of 3.4 Å. The effects of RNA atoms outside the quantum region, as well as water molecules and ions in the solvent, were represented as point charges uniformly distributed on the molecular surface of the quantum region and determined from Poisson-Boltzmann calculations using the solinprot program in the MEAD package^72,73^. The quantum region was assigned a local dielectric constant of 1, the remaining RNA region a dielectric constant of 4, and the solvent region a dielectric constant of 80.

Within the AF-QM/MM framework, quantum-mechanical shielding calculations were performed with ORCA 6.1.1 using the same settings as Roy et al^14,74^. Specifically, shielding tensors were computed using the GIAO-DFT formalism with the OLYP functional and the pcSseg-1 basis set optimized for chemical shifts^75,76^. Reference shieldings were computed for tetramethylsilane (TMS) using the same level of theory and used to convert the isotropic components of the calculated shielding tensors into predicted chemical shifts. Predicted shifts were averaged over all conformers in the ensemble, and resonance-type-specific linear corrections were applied to the ensemble-averaged values before comparison with experiment^14,30,77^.

Apart from looking at the specific atom type chemical shift calculation accuracy, we also used N-weighted RMSD and N-weighted R² to evaluate the overall agreement between predicted and experimental chemical shifts across atom types. For each atom type (*k*), the RMSD and R² values were calculated separately, and the overall metrics were then obtained by weighting each atom type according to the number of assigned data points.

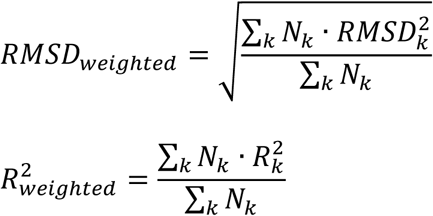

Here, *N_k_* denotes the number of chemical shift data points for atom type *k*. This weighting scheme gives proportionally greater contribution to atom types with more experimental assignments and provides a global measure of chemical shift prediction accuracy.

### Ensemble-based virtual screening

Ensemble-based virtual screening (EBVS) was performed using Glide in the Schrödinger Suite with RNA ensembles containing 50 conformers^78,79^. The HIV-1 TAR small-molecule library was taken from Ganser et al^42^. Each RNA conformer was prepared with the Protein Preparation Wizard in Schrödinger, and the ligand library was processed with LigPrep, with protonation and ionization states generated by Epik at pH 7.4. For each conformer, a docking grid with a box size of 40 × 40 × 40 Å was centered on the mass center of the full RNA. Docking was carried out in Glide standard precision using the RNA-specific scoring function, and each ligand was docked independently against all 50 conformers^80^. The final score for each ligand was calculated as a Boltzmann-weighted average over the full ensemble, in which each conformer contributed according to its relative Boltzmann population and the weighted docking scores were summed across all conformers, following the procedure described previously^42^.

Virtual-screening performance was evaluated using receiver operating characteristic (ROC) curves, which plot the true-positive rate against the false-positive rate across a range of docking-score thresholds. Overall discrimination was summarized by the area under the ROC curve (AUC), with values of 0.5 and 1.0 corresponding to random ranking and perfect separation, respectively.

### Mahalanobis distance

To quantify ensemble coverage of experimental RNA-ligand complex structures, we computed the Mahalanobis distance in interhelical Euler-angle space^41^. Mahalanobis distance measures how far a point lies from the center of a distribution while accounting for correlations through the covariance matrix. For the inter-helical Euler angles, Mahalanobis distance was calculated as:

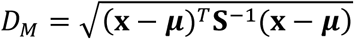

where **x** is the experimental structure’s Euler angle vector, ***μ*** is the ensemble mean, and **S** is the covariance matrix.

### Geodesic Rotation Distance

Following the approach used previously to compare simulated HIV-1 TAR interhelical conformations with ligand-bound structures, the interhelical Euler angles were converted to rotation matrices, and the difference between two orientations was quantified as the magnitude of their relative single-axis rotation^81^. This quantity is equivalent to the geodesic rotation distance and was calculated as

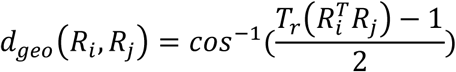

For each ensemble and ligand-bound PDB structure, the minimum distance between any ensemble conformer and any experimental model was reported. Direct coverage was additionally quantified as the fraction of ensemble conformers within 20° of at least one experimental model, consistent with the rotational resolution previously used to assess sampling of TAR interhelical orientations.

### Trajectory analysis and structure visualization

Trajectory analysis was carried out using CPPTRAJ^82^. All 3D structural images presented in the manuscript were generated with PyMOL^83^.

## Data Availability Statement

The ensemble files generated from the different enhanced-sampling MD simulations, together with the analysis code used to generate the figures in this manuscript, are available at https://github.com/Ken-Lab-ScrippsResearch/Li-Ken-RNA-ESMD.

## Supporting Information

RDC correlation plots for TAR ensembles generated using T-REMD temperature windows of 298 to 330 K and 298 to 380 K are shown in Figure S1. Atom-type-resolved QM/MM chemical-shift validation of the TAR ensembles is shown in Figure S2. ROC AUC curves for ensemble-based virtual screening using TAR ensembles generated by cMD, GaMD, Rex-GaMD, and REST2 are shown in Figure S3. RDC correlation plots and three-dimensional ensembles of the preQ1 class I riboswitch generated using GaMD, Rex-GaMD, and REST2 are shown in Figure S4. RDC correlation plots and three-dimensional ensembles of the UUCG tetraloop generated using GaMD, Rex-GaMD, and REST2 are shown in Figure S5. Computational costs for the HIV-1 TAR simulations, reported as aggregate GPU-hours per 1000 ns trajectory, are shown in Figure S6.

## Acknowledgments

We thank Prof. Hashim M. Al-Hashimi of Columbia University, Honglue Shi of the University of California, Berkeley, and Manuel Llanos of Scripps Research, for providing helpful suggestions for this work. This work was supported by startup funds from Scripps Research, the Center for Structural Biology of HIV RNA (CRNA) (U54 AI170660-01), and the NIH Director’s Early Independence Award (DP5OD037420). This work used the Delta and DeltaAI systems at the National Center for Supercomputing Applications through allocation BIO250257 from the Advanced Cyberinfrastructure Coordination Ecosystem: Services & Support program, which is supported by National Science Foundation grants 2138259, 2138286, 2138307, 2137603, and 2138296. Additional computational resources were provided by the high-performance computing facilities at Scripps Research.

