## Supplemental Information for "Enhanced-Sampling Molecular Dynamics Recovers Rare Functional RNA Conformations Across Diverse Structural Contexts"

Deng Li[1,2] and Megan Ken[1,2*]

[1]Department of Integrative Structural and Computational Biology, Scripps Research, La Jolla, CA 92037

[2]Department of Immunology and Microbiology, Scripps Research, La Jolla, CA 92037

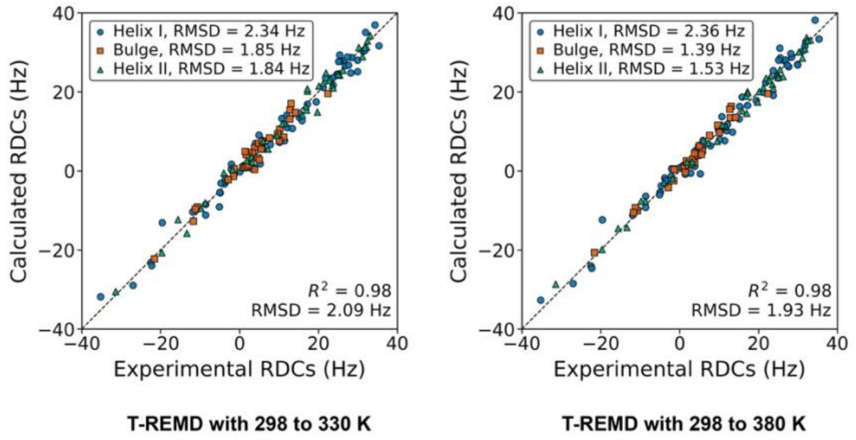


**Figure S1.** Effect of T-REMD temperature window on TAR RDC-refined ensemble accuracy. RDC correlation plots for TAR ensembles generated using narrower T-REMD temperature windows of 298 to 330 K and 298 to 380 K.

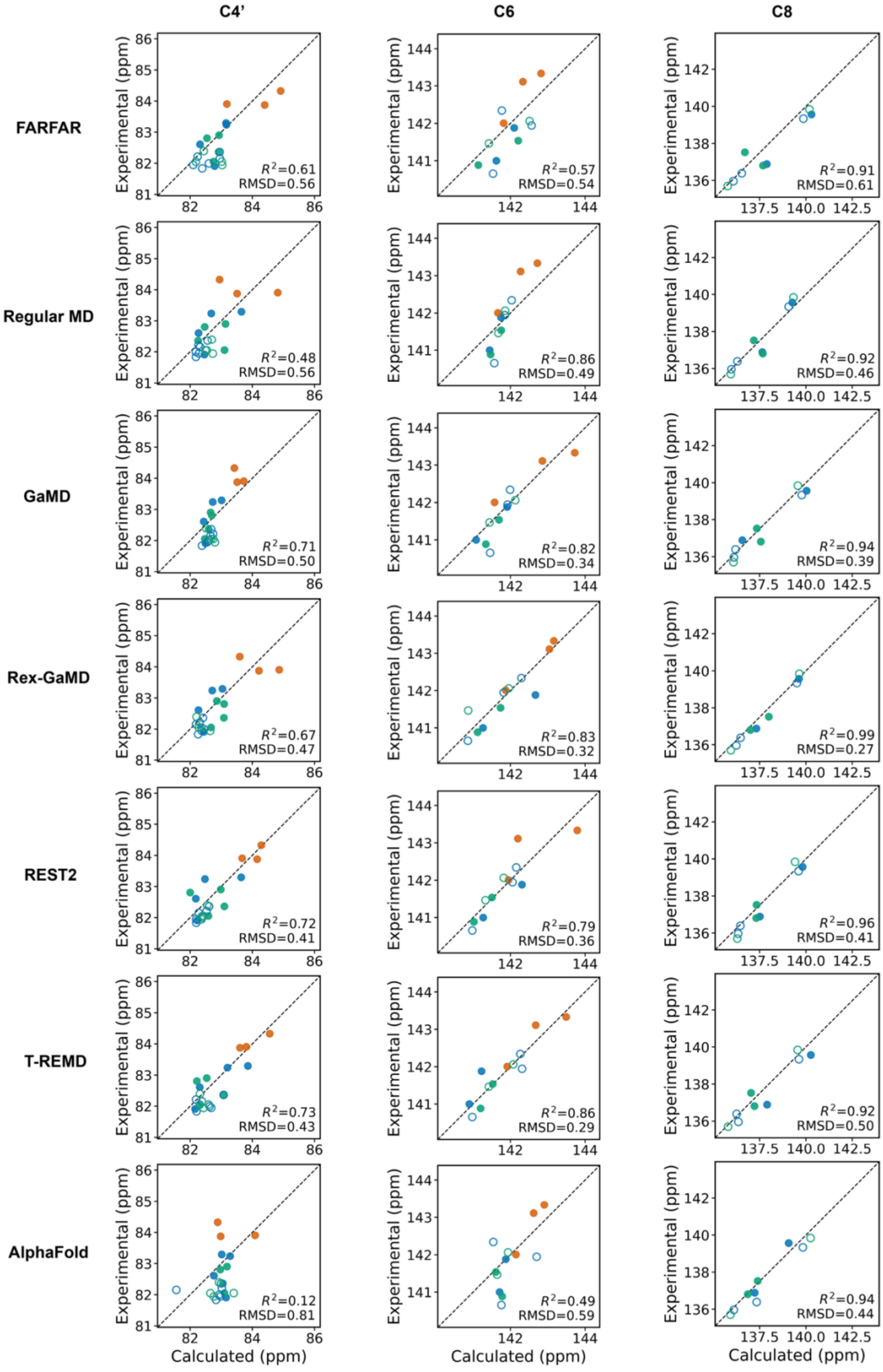

C4’
C6
C8
FARFAR
$R^2$=0.61 RMSD=0.56
$R^2$=0.57 RMSD=0.54
$R^2$=0.91 RMSD=0.61
Regular MD
$R^2$=0.48 RMSD=0.56
$R^2$=0.86 RMSD=0.49
$R^2$=0.92 RMSD=0.46
GaMD
$R^2$=0.71 RMSD=0.50
$R^2$=0.82 RMSD=0.34
$R^2$=0.94 RMSD=0.39
Rex-GaMD
$R^2$=0.67 RMSD=0.47
$R^2$=0.83 RMSD=0.32
$R^2$=0.99 RMSD=0.27
REST2
$R^2$=0.72 RMSD=0.41
$R^2$=0.79 RMSD=0.36
$R^2$=0.96 RMSD=0.41
T-REMD
$R^2$=0.73 RMSD=0.43
$R^2$=0.86 RMSD=0.29
$R^2$=0.92 RMSD=0.50
AlphaFold
$R^2$=0.12 RMSD=0.81
$R^2$=0.49 RMSD=0.59
$R^2$=0.94 RMSD=0.44
Experimental (ppm)
Calculated (ppm)

**Figure S2.** Atom-type-resolved QM/MM chemical shift validation of TAR ensembles. Correlations between experimental and ensemble-averaged QM/MM-predicted chemical shifts for C4′, C6, and C8 atoms in TAR ensembles generated from FARFAR, conventional MD, GaMD, Rex-GaMD, REST2, T-REMD, and AlphaFold 3 conformational libraries.

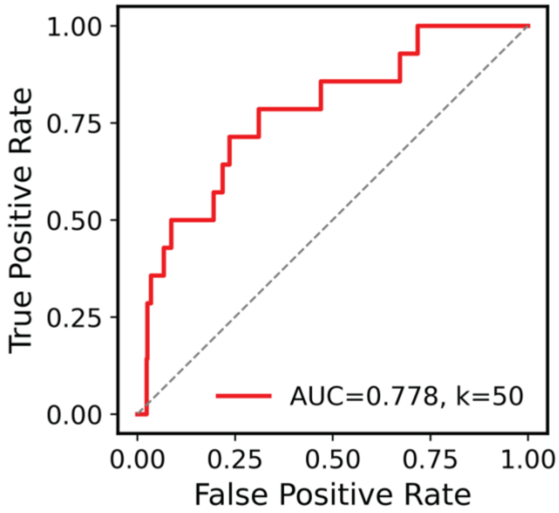


AUC curve using ensemble from conventional MD

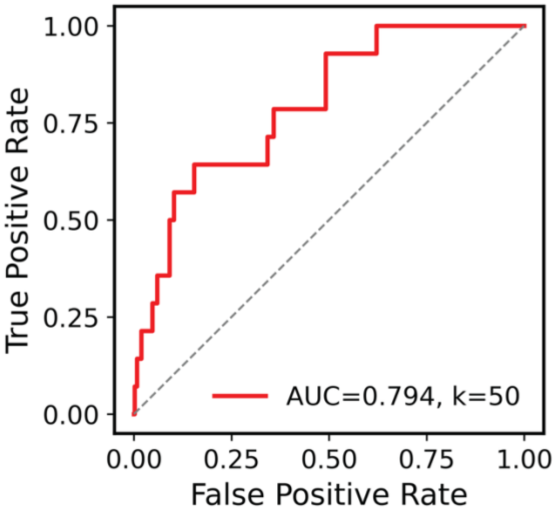


AUC curve using ensemble from GaMD

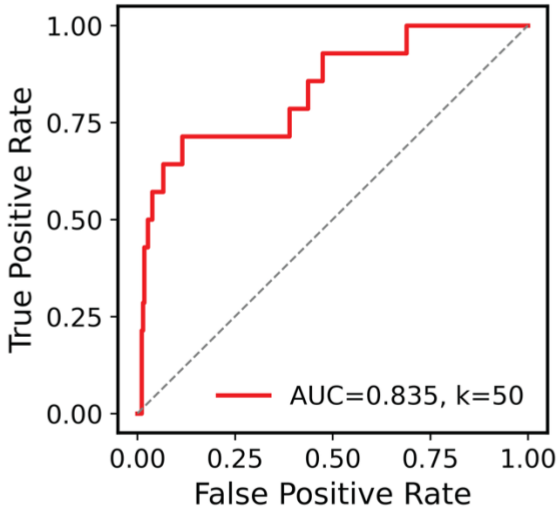


AUC curve using ensemble from Rex-GaMD

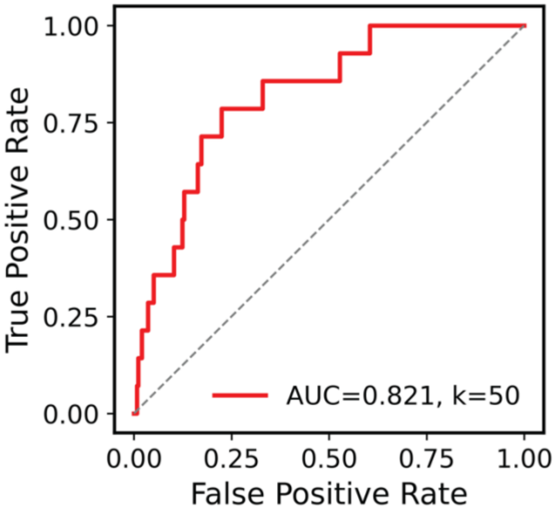


AUC curve using ensemble from REST2

**Figure S3.** Ensemble-based virtual screening performance for MD-derived TAR ensembles. ROC AUC curve for ensemble-based virtual screening using TAR ensembles generated from cMD, GaMD, Rex-GaMD, and REST2 conformational libraries.

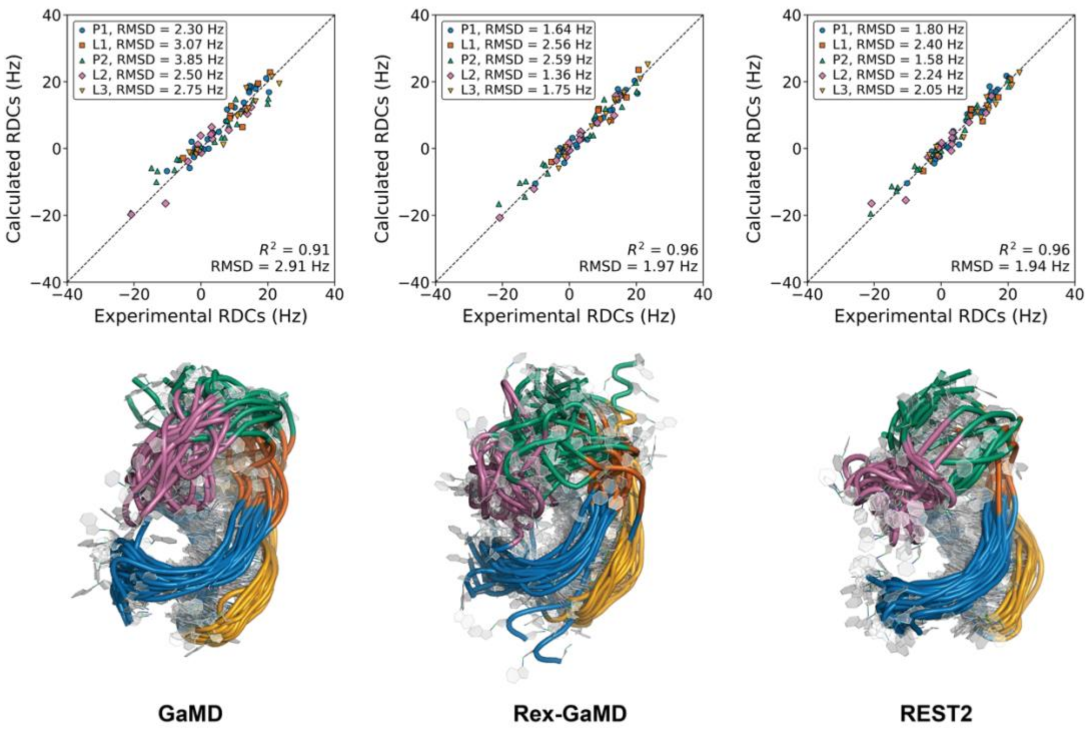


**Figure S4.** Enhanced-sampling ensemble construction for the preQ1 class I riboswitch. RDC correlation plots and 3D ensemble for preQ1 riboswitch generated using GaMD, Rex-GaMD, and REST2.

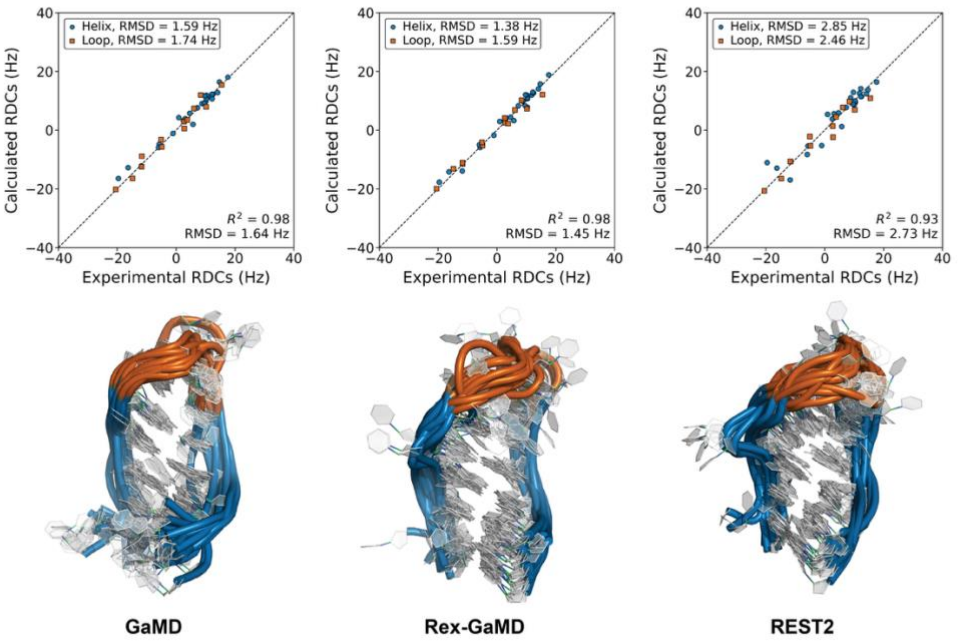


**Figure S5.** Enhanced-sampling ensemble construction for the UUCG tetraloop. RDC correlation plots and 3D ensemble for UUCG tetraloop generated using GaMD, Rex-GaMD, and REST2.

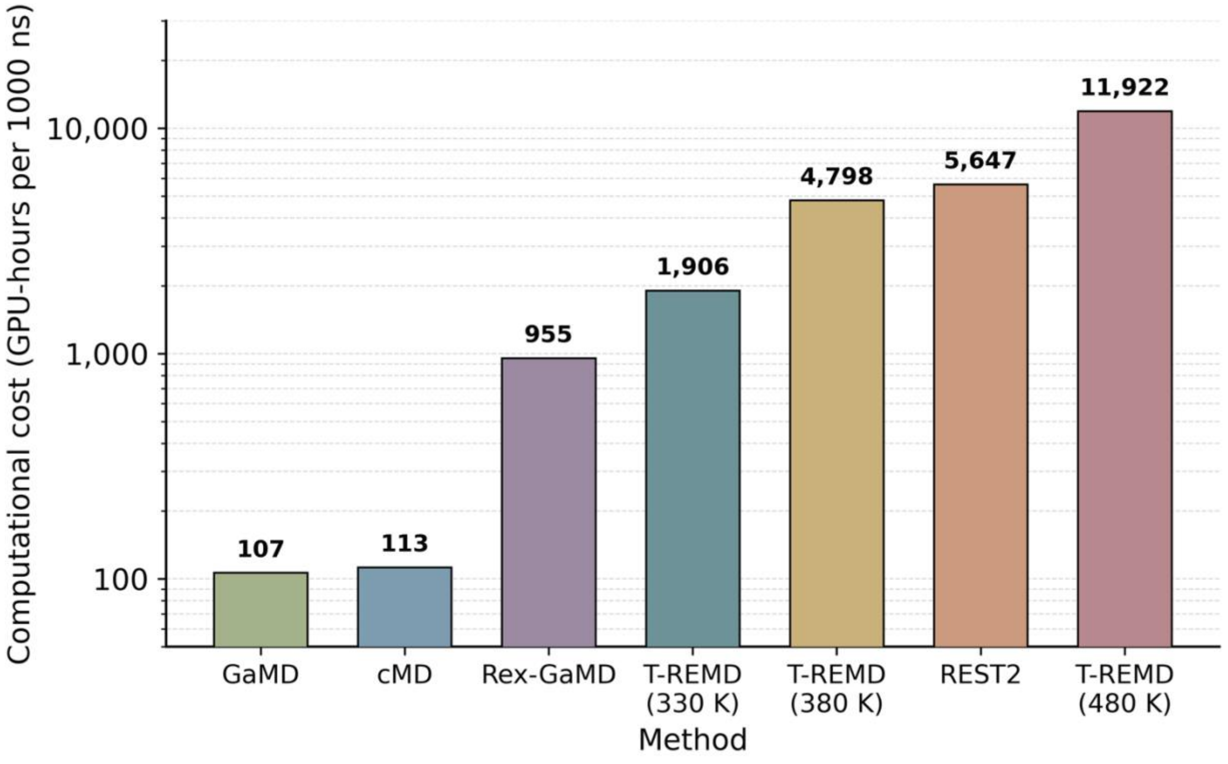


**Figure S6.** Computational cost (GPU-hours per 1000 ns trajectory) for each method (HIV-1 TAR), benchmarked on NVIDIA RTX A5000 (24 GB) GPUs. Cost is the aggregate cost across all GPUs used.
